# Comparison of SARS-CoV-2 spike protein binding to human, pet, farm animals, and putative intermediate hosts ACE2 and ACE2 receptors

**DOI:** 10.1101/2020.05.08.084061

**Authors:** Xiaofeng Zhai, Jiumeng Sun, Ziqing Yan, Jie Zhang, Jin Zhao, Zongzheng Zhao, Qi Gao, Wan-Ting He, Michael Veit, Shuo Su

**Author notes:** co-first author (Xiaofeng Zhai, Jiumeng Sun). co-last senior author (Michael Veit, Shuo Su).

## Abstract

The emergence of a novel coronavirus, SARS-CoV-2, resulted in a pandemic. Here, we used recently released X-ray structures of human ACE2 bound to the receptor-binding domain (RBD) of the spike protein (S) from SARS-CoV-2 to predict its binding to ACE2 proteins from different animals, including pets, farm animals, and putative intermediate hosts of SARS-CoV-2. Comparing the interaction sites of ACE2 proteins known to serve or not serve as receptor allows to define residues important for binding. From the 20 amino acids in ACE2 that contact S up to seven can be replaced and ACE2 can still function as the SARS-CoV-2 receptor. These variable amino acids are clustered at certain positions, mostly at the periphery of the binding site, while changes of the invariable residues prevent S-binding or infection of the respective animal. Some ACE2 proteins even tolerate the loss or the acquisition of N-glycosylation sites located near the S-interface. Of note, pigs and dogs which are not or not effectively infected, respectively, have only a few changes in the binding site have relatively low levels of ACE2 in the respiratory tract. Comparison of the RBD of S of SARS-CoV-2 with viruses from bat and pangolin revealed that the latter contains only one substitution, whereas the bat virus exhibits five. However, ACE2 of pangolin exhibit seven changes relative to human ACE2, a similar number of substitutions is present in ACE2 of bats, raccoon, and civet suggesting that SARS-CoV-2 may not especially adapted to ACE2 of any of its putative intermediate hosts. These analyses provide new insight into the receptor usage and animal source/origin of SARS-COV-2.

**IMPORTANCE:** SARS-CoV-2 is threatening people worldwide and there are no drugs or vaccines available to mitigate its spread. The origin of the virus is still unclear and whether pets and livestock can be infected and transmit SARS-CoV-2 are important and unknown scientific questions. Effective binding to the host receptor ACE2 is the first prerequisite for infection of cells and determines the host range. Our analysis provides a framework for the prediction of potential hosts of SARS-CoV-2. We found that ACE2 from species known to support SARS-CoV-2 infection tolerate many amino acid changes indicating that the species barrier might be low. However, the lower expression of ACE2 in the upper respiratory tract of some pets and livestock means more research and monitoring should be done to explore the animal source of infection and the risk of potential cross-species transmission. Finally, the analysis also showed that SARS-CoV-2 may not specifically adapted to any of its putative intermediate hosts.

## INTRODUCTION

As of the 30^th^ of April 2020, the ongoing pandemic of a novel coronavirus, severe acute respiratory syndrome coronavirus 2 (SARS-CoV-2), has developed into a global challenge with the number of total confirmed cases exceeding 3 million including more than 200 thousands fatalities, thereby causing a major loss to global public health and the world economy. This disease is referred to as the 2019 coronavirus disease (COVID-19) by World Health Organization (WHO) and was defined as a public health emergency of international concern (PHEIC) on the 30^th^ of January 2020. Its main clinical symptoms include fever, fatigue, and dry cough. A rather large proportion of patients become critically ill with acute respiratory distress syndrome, similar to patients with severe acute respiratory syndrome (SARS) caused by SARS coronavirus (SARS-CoV) (1–3). SARS emerged in China in 2002-2003 and also rapidly spread worldwide but was contained by public health measures. It is thought that bats and palm civets are the natural and intermediate reservoirs of SARS-CoV (4, 5). Likewise, research suggests that SARS-CoV-2 might have originated also from bats and that pangolins might be the potential intermediate host (6–9). Specifically, SARS-CoV-2 has a high nucleotide sequence identity with Bat-CoV-RaTG13-like virus except for the middle part of its genome encoding the spike protein which might have derived via recombination from a Pangolin-CoV-like virus (6, 7, 10–12). A previous study showed that SARS-CoV-2 replicates poorly in dogs, pigs, but cats are permissive to infection (13). However, whether pets can become new hosts of SARS-CoV-2 needs to be clarified further.

The structure of the trimeric spike protein (S) of SARS-CoV-2, the major factor that determines cell and host tropism, has been determined (14–17). It is cleaved by host proteases into the S1 subunit, which contains the receptor binding domain (RBD), and S2, which mediates fusion of the virion with cellular membranes (18). Proteolytical cleavage of S at two sites (S1/S2 and S2’) is required to prime the protein to execute its fusion activity. Cleavage is performed by the serine transmembrane protease TMPRSS2, but in its absence lysosomal cathepsin proteases B and L can substitute (19). S of SARS-CoV-2 contains an insertion of four amino acids that creates a cleavage site for the ubiquitous protease furin (16, 20, 21).

Angiotensin-converting enzyme 2 (ACE2) belongs to the angiotensin-converting enzyme family of dipeptidyl carboxyl dipeptidase and has considerable homology with human angiotensin-converting enzyme 1. SARS-CoV-2 and SARS-CoV share the same membrane protein, angiotensin converting enzyme 2 (ACE2), as the cellular receptor (16, 22). In this study, we used a comparative bioinformatics and structural approach to better understand the source and host range of emerging SARS-CoV-2. We compared recently released X-ray structures of human ACE2 bound to the receptor-binding domain of S from SARS-CoV-2 with the structure of S of SARS-CoV bound to human ACE2 (23–26). Using ACE2 sequences of species that can serve and not serve as a receptor for SARS-CoV-2 we propose amino acids crucial for binding (12). We report that pets (cats) and domestic animals are at risk of infection with SARS-CoV-2 since they have fewer amino acid changes at the ACE2-S interface compared with ACE2 proteins from animals that are known to serve as the SARS-CoV-2 receptor. In China, Europe, and other countries, pets have long accompanied humans and thus they may be a neglected source worthy of further research. In addition, using RBD sequences of putative intermediate hosts, we consider which of them is more likely to bind to human ACE2. We also compared O-and N-glycosylation sites and a putative integrin binding motif in the S proteins of these viruses.

## RESULTS AND DISCUSSION

### Interacting amino acids of human ACE2 and SARS-CoV-2 and SARS-CoV S proteins

Three crystal structures of the receptor-binding domain of S from SARS-CoV-2 in a complex with its receptor, human ACE2, has been resolved recently (24–26). The protein interaction surface (pdb-file 6MOJ) is depicted in Fig. 1A (26). A total of 17 residues in the spike protein are in contact with 20 amino acids in ACE2, eight of which (labelled in orange) form hydrogen bonds with 13 residues in S (see also Table 1). The other interactions are mostly hydrophobic involving many tyrosine residues in the viral spike. A substantial part of the binding energy might be provided by the formation of a salt bridge between a negatively charged Asp30 in ACE2 with a positively charged Lys417 in the spike protein, which is located in the middle of the interaction surface.

**Table 1.**
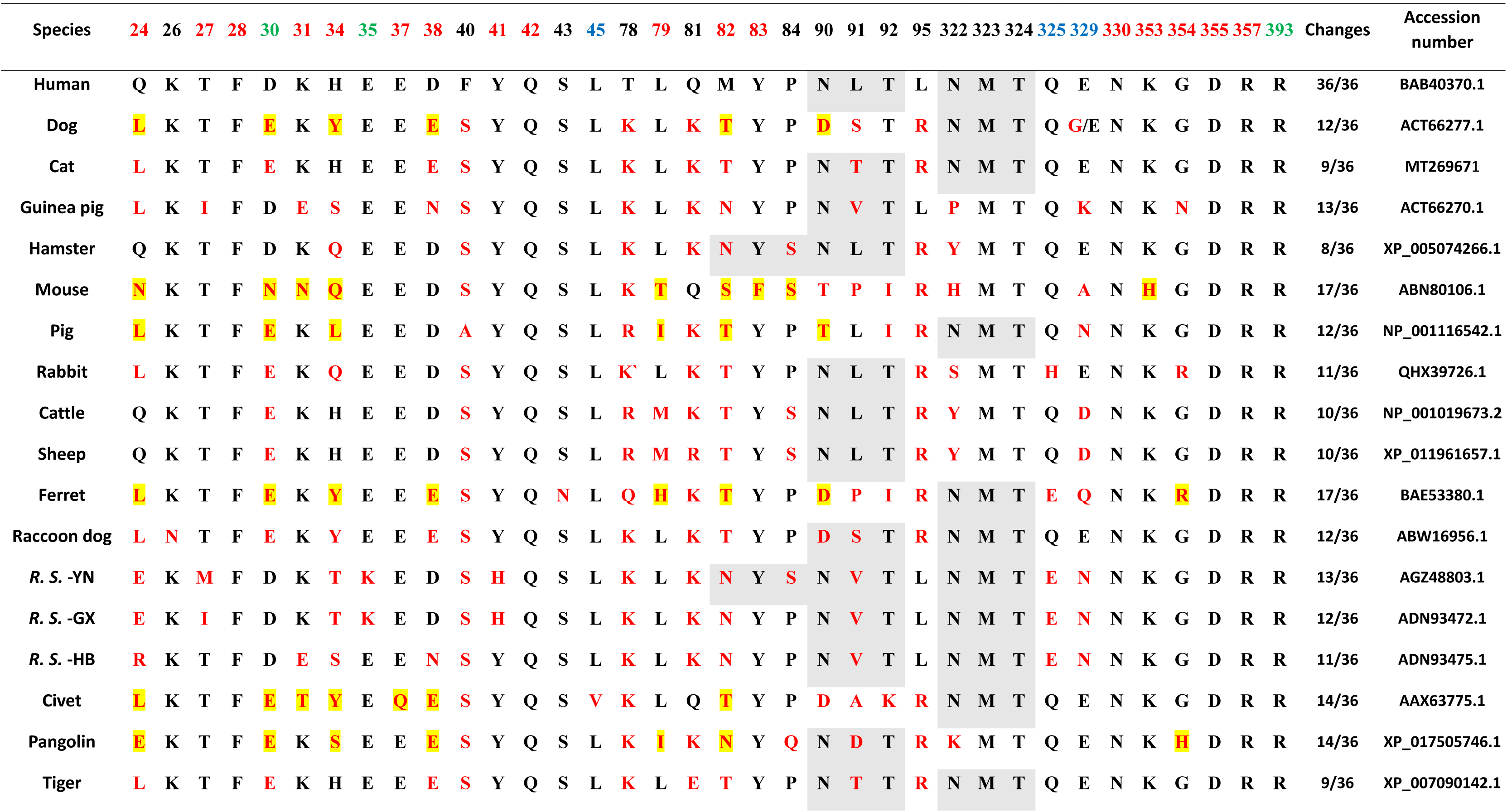
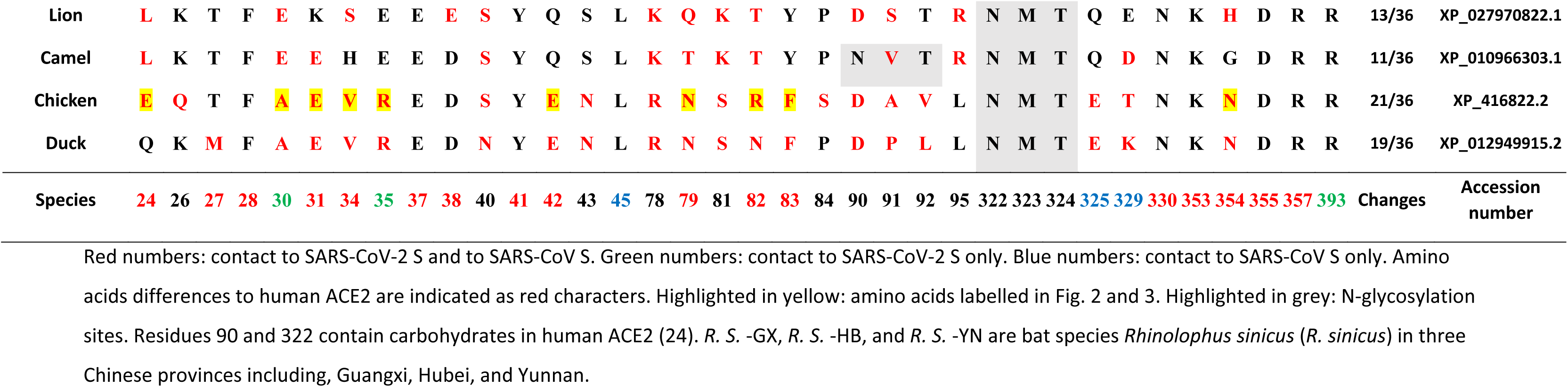
Comparison of the ACE2 residues interfacing with SARS-CoV and with SARS-CoV-2 receptor-binding domain (RBD) of different species

**Fig. 1.**
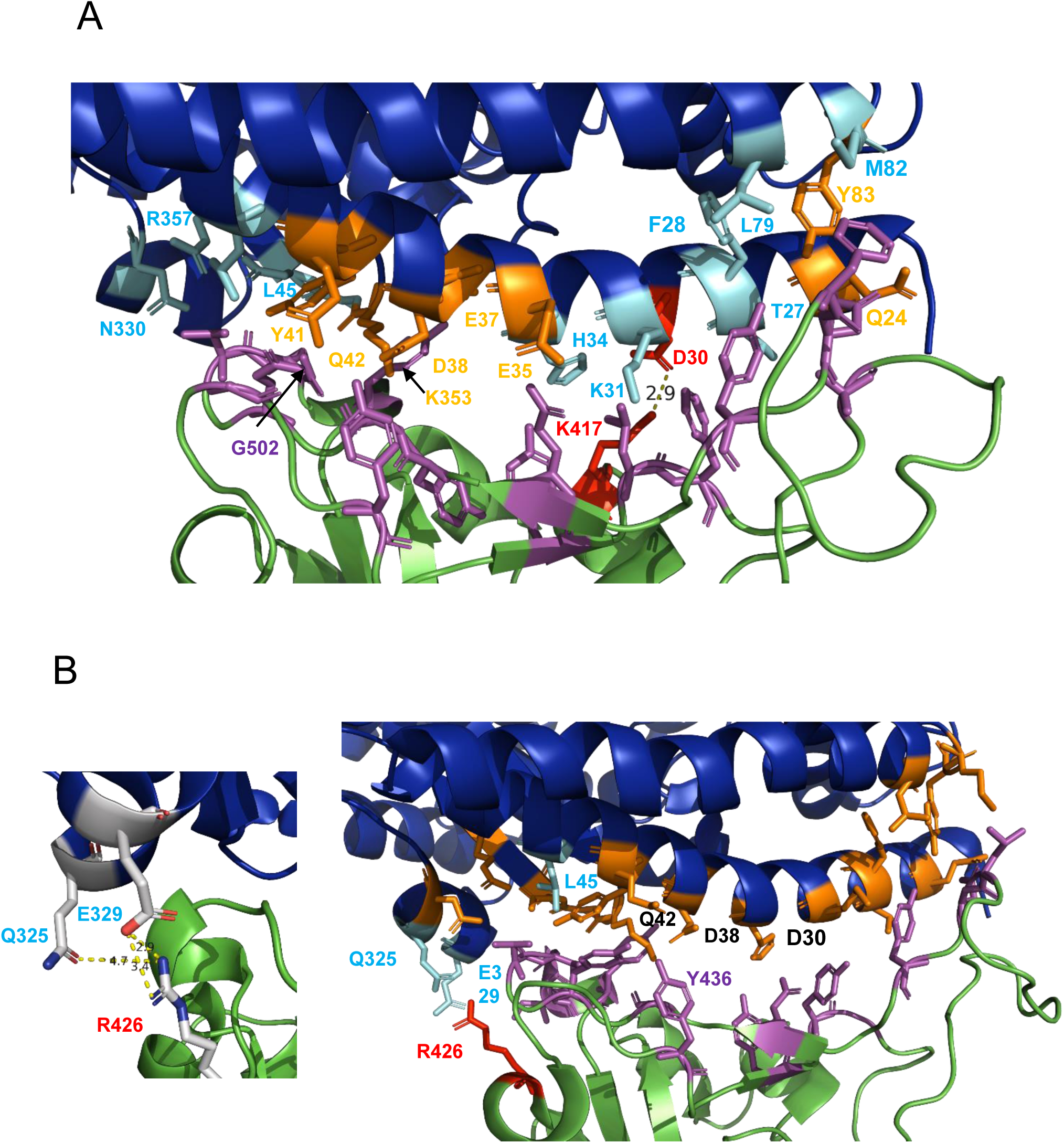
(A) Interaction surface of SARS-CoV-2 S (green) and human ACE2 (blue). Amino acids involved in contact are shown in magenta (S) or in ACE2 in orange (amino acids forming hydrogen bonds) and cyan (hydrophobic interaction) sticks. D30 forms a salt bridge with K417 in S of SARS-CoV-2. The figure was created with PyMol from pdb file 6MOJ. (B) Interaction surface between SARS-CoV S (green) and human ACE2 (blue). Amino acids making the contacts are shown in magenta (S) or orange (ACE2) sticks. The three amino acids in ACE2 that are not involved in binding of S from SARS-CoV-2 are shown in cyan sticks. The side chain of D30 that forms a salt bridge with S of SARS-CoV-2 is pointing away from the interacting surface. Inset: Detail of the interaction between R426 that forms a salt bridge with E329 and a hydrogen bond with Q325. The figure was created with PyMol from pdb file 2AJF.

A second crystal structure (pdb file 6LZG) identified one more contact site: Ser19 in ACE2 interacts with Ala457 and Gly458. Ser19 is the first amino acid in ACE2 of all crystals and hence its side chain might be more flexible (24). The third crystal structure (pdb-file 6VW1) identified three more sites, Leu45, Gln325, and Glu329, that were identified in the two other papers to be unique for binding to S of SARS-CoV, but not the salt bridge involving Asp30 (25). However, the authors used a chimeric S protein that contains the receptor-binding domain of SARS-CoV plus the loop that maintains a salt bridge between Arg426 with Glu329 in ACE2 (see below). Thus, this chimeric spike might not display the authentic binding properties of SARS-CoV-2.

It was reported in some studies that S of SARS-CoV-2 binds with higher affinity to its receptor than S of SARS-CoV (14, 16, 24, 27). Wang et al. (2020) and Lan et al. (2020) identified more amino acids in SARS-CoV-2 compared with SARS-CoV interacting with ACE2 that also form more hydrogen bonds and van-der-Waal contacts (24, 26). To visualize the amino acids involved in binding of both S proteins we used the crystal structure of S from SARS-CoV bound to human ACE2 (23). The contact surface of S is formed by a similar number of residues (16) which interact with 20 amino acids in ACE2, but three of the latter residues are unique for binding to S of SARS-CoV (Fig. 1B, labelled in cyan, and Table 1). Surprisingly, two of the unique residues (Gln325, Glu329) are involved in the formation of the only salt bridge with Arg426 in S, which is located at the periphery of the binding site. The side chain of Asp30, that forms the salt bridge with SARS-CoV-2 S, is pointing away from the interacting surface and is thus not available for binding. In summary, although most of residues in ACE2 involved in binding to S of SARS-CoV-2 and SARS-CoV are identical, some are unique suggesting that coronaviruses can adapt in multiple ways to the human ACE2 receptor.

### Comparison of the interacting surface of ACE2 proteins that serve and not serve as SARS-CoV-2 receptor

It has been shown that transfection of HeLa cells with genes encoding the human, pig, civet, and bat ACE2 receptor makes them susceptible to infection with SARS-CoV-2, but not with ACE2 from mice (12). To estimate which of the interacting amino acids in ACE2 are essential for binding of S, we compared the ACE2 sequences from humans with that from the other species. Table 1 shows the amino acids that make contacts with S from both SARS-CoV-2 and SARS-CoV (red numbers), only with S from SARS-CoV (blue numbers), only with S from SARS-CoV-2 (green numbers) as well as some variable amino acids in the vicinity (black numbers), and some encoding N-glycosylation sites (highlighted in grey).

**Pig ACE2** contains five amino acid substitutions in the interacting surface with S from SARS-CoV-2 relative to human ACE2 (Fig. 2A). Three of the exchanges are located at the periphery of the binding site. Leu79 and Met82, which interact with Phe486 in S, are conservatively substituted by Ile and Thr, respectively. Gln24, which forms a hydrogen bond to N487 in S, is replaced by Leu, which cannot form hydrogen bonds. His34 in the centre of the binding site is also substituted by a Leu residue, which is larger, but cannot from a hydrogen bond. Furthermore, Asp30, which forms the central salt bridge, is exchanged by a Glu residue. Since Glu is also negatively charged, the salt bridge to Lys417 most likely remains intact. It might even become stronger since the side chain of Glu is larger and hence the distance between negatively and positively charged residues becomes smaller. In addition, the N-glycosylation site Asn90 in human ACE2 is destroyed by an exchange to Thr.

**Fig. 2.**
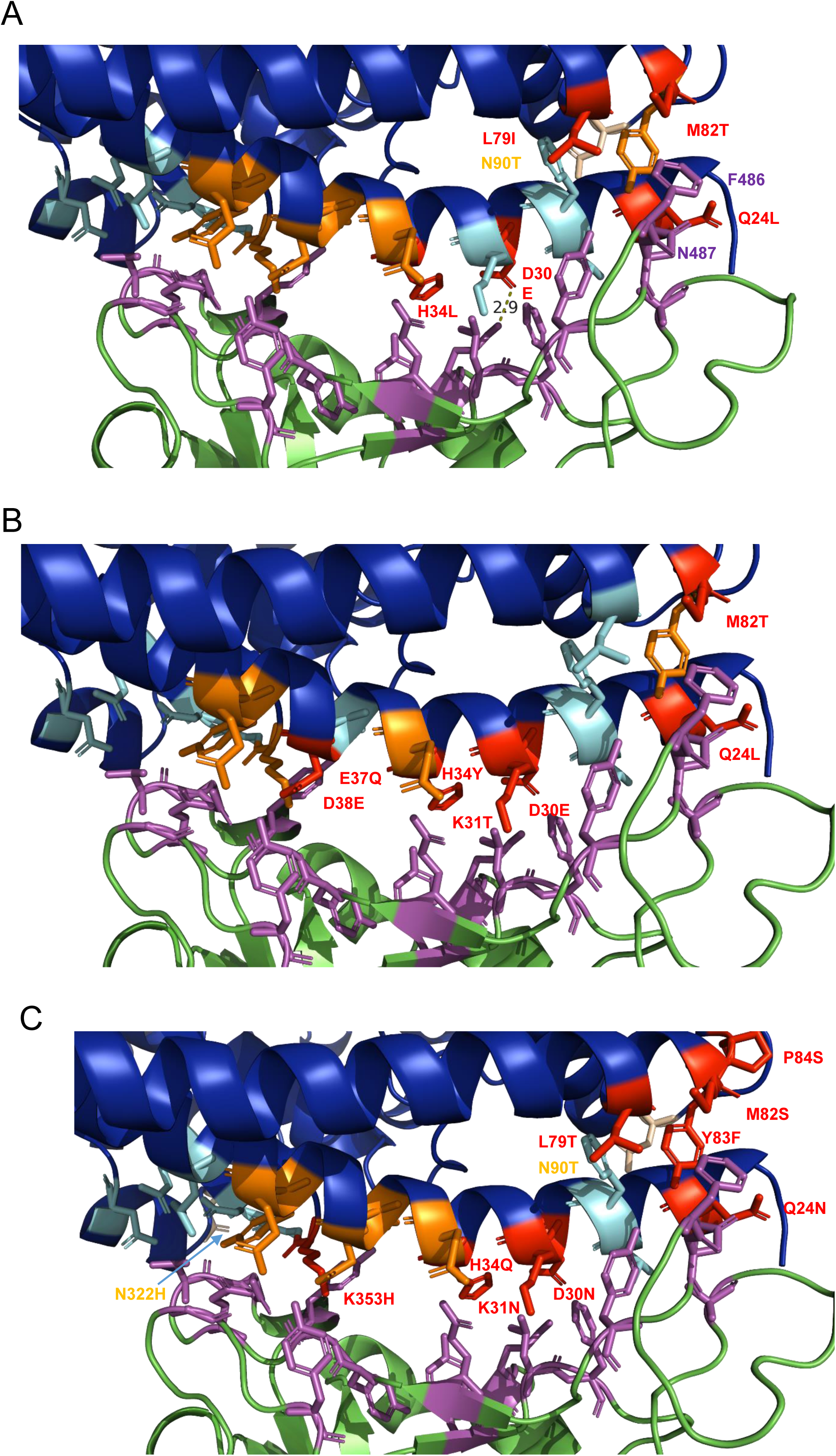
(A) Amino acid changes of human ACE2 and pigs ACE2. Amino acid changes in ACE2 from pigs compared with human ACE2 are highlighted in red. N90 is not an interaction site, but the change N90T destroys the N-glycosylation site present in human ACE2. (B) Amino acid changes of human ACE2 and civet ACE2. Amino acid changes in ACE2 from civet compared with human ACE2 are highlighted in red. (C) Amino acid changes of human and mouse ACE2. Amino acid changes in human ACE2 compared with mouse ACE2 are highlighted in red. P84 in human ACE2 does not interact with amino acids in S of SARS-CoV-2 but might affect the secondary structure. N90T and N322H destroy the N-glycosylation site in human ACE2.

Three amino acid sequences of the ACE2 gene from the **bat** species *Rhinolophus sinicus* (*R. sinicus*) are present in the database. The bats were sampled from *R. sinicus* colonies in three Chinese provinces including, Guangxi (*R. S.* -GX), Hubei (*R. S.* -HB), and Yunnan (*R. S.* -YN) (28, 29). Strikingly, although their overall amino acid identity is very high (99%), they exhibit large amino acid differences in the N-terminal amino acids that contact the S protein (Table 1). Since the accession number of the bat ACE2 is not specified in the publication that demonstrates that it confers susceptibility to SARS-CoV-2 infection, it is presently unclear which of them is recognized by the viral spike protein (12). In any case, bat ACE2 proteins contain at least five changes relative to human ACE2. Three of them are substituted in all bat ACE2 sequences and involve the same residues as in pig ACE2, but they are replaced by other amino acids. Gln24 is replaced by the negatively charged Glu or by a positively charged Arg; His34 is substituted by Thr or Ser; and Met82 is substituted by an Asn. In one bat sequence (*R. S.* -YN) the next but one Pro residue 84 is also replaced by a Ser, which creates a new N-glycosylation site (NXS) at position 82. In addition, in two bat sequences (*R. S.* -YN and *R. S.* -GX) Thr27 is substituted by Met or Iso and Tyr41, which forms a hydrogen bond with Thr500 in S, is replaced by the smaller His residue. The most drastic substitution in these two bat ACE2 sequences is Glu35, which forms a hydrogen bond with Gln493 in S, by a positively charged Lys residue. The ACE2 sequence obtained from a bat in the Hubei province, which exhibits the most amino acid substitutions relative to the two other ones, does not contain the latter three changes, but instead Arg31 is replaced by a negatively charged Glu and Asp38 by Asn. The reason why bats exhibit so many changes in residues that interact with S is striking and requires further investigation. However, it is tempting to speculate that local co-evolution between bats and coronaviruses drive these amino acid changes.

To get further insight into the amino acids not essential for binding to S, we analysed the ACE2 from civets (*Paguma larvata*) that has been shown to serve as receptor for SARS-CoV-2 (12). **Civet ACE2** contains seven amino acid changes relative to human ACE2 (Fig. 2B). Three of them (Gln24Leu, Asp30Glu, and Met82Thr) are identical to the substitutions in pig ACE2. Another (His34) is at the same position, but exchanged to a different amino acid, Tyr instead of Leu. Another unique, but conservative substitution, Leu45V, is located at the periphery of the binding site, whereas the other three are in the centre. Asp38Glu is a conservative change, but K31T and E37Q replace a charged by an uncharged amino acid.

**Mouse ACE2**, which does not support infection of cells with SARS-CoV-2, has eight amino acid substitutions in the interacting surface with S of SARS-CoV-2 (Fig. 2C). Three of the sites, Gln24, His34, and Met82 are also replaced in the ACE2 proteins from the two other species and are thus unlikely to be the decisive elements that prevent binding. Leu79 interacts with Phe486 in S (Fig. 6A) and is exchanged by a Thr. In contrast to bat ACE2, the substitution at position 82 does not create a N-glycosylation site in mouse ACE2 since it is exchanged to Ser. Note also that the two used N-glycosylation sites near the interacting surface in human ACE2, Asn322, and Asn90, are lost in mouse ACE2 due to exchanges of the Asn residues. The other four exchanges Asp30Asn, Tyr83Phe, Lys31Asn, and Lys353His are more important for preventing binding to mouse ACE2 as discussed in more detail below.

In summary, binding of S of SARS-CoV-2 to ACE2 proteins tolerates a surprisingly large number of amino acid changes in the interaction surface, five in pig ACE2 and seven in civet ACE2. Even the acquisition of an additional N-glycosylation site at position 82 due to two substitutions in bat ACE2 (M82N, P84S) does not prevent SARS-CoV-2 to use bat ACE2 as receptor in transfected cells. This is in contrast to binding of S of SARS-CoV to rat ACE2, where a glycan attached to the same position prevents binding (23).

### Comparison of the interacting surface of human ACE2 and ACE2 from pet animals

Next, we asked whether receptor binding might present a species barrier for infection of pets with SARS-CoV-2. **Dog ACE2** contains five amino acid changes in the amino acids in contact with S (Fig. 3A). Residues 24, 30, 34, and 82 are also replaced in pig ACE2, even to the same amino acid at four of the sites. A unique change in dog ACE2 is Asp38Glu, but since this is a conservative change it is unlikely to affect the binding properties of S. Like in pigs, dog ACE2 lacks the glycosylation site at position 90. Asn90 is replaced by Asp in dogs, but to Thr in pigs. Note that the sequence of dog ACE2 in the database contains a non-conservative exchange at position 329 (Glu to Gly), which contacts the S protein of SARS-CoV. This exchange is not present in the ACE2 from a beagle that we sequenced (accession number MT269670).

**Fig. 3.**
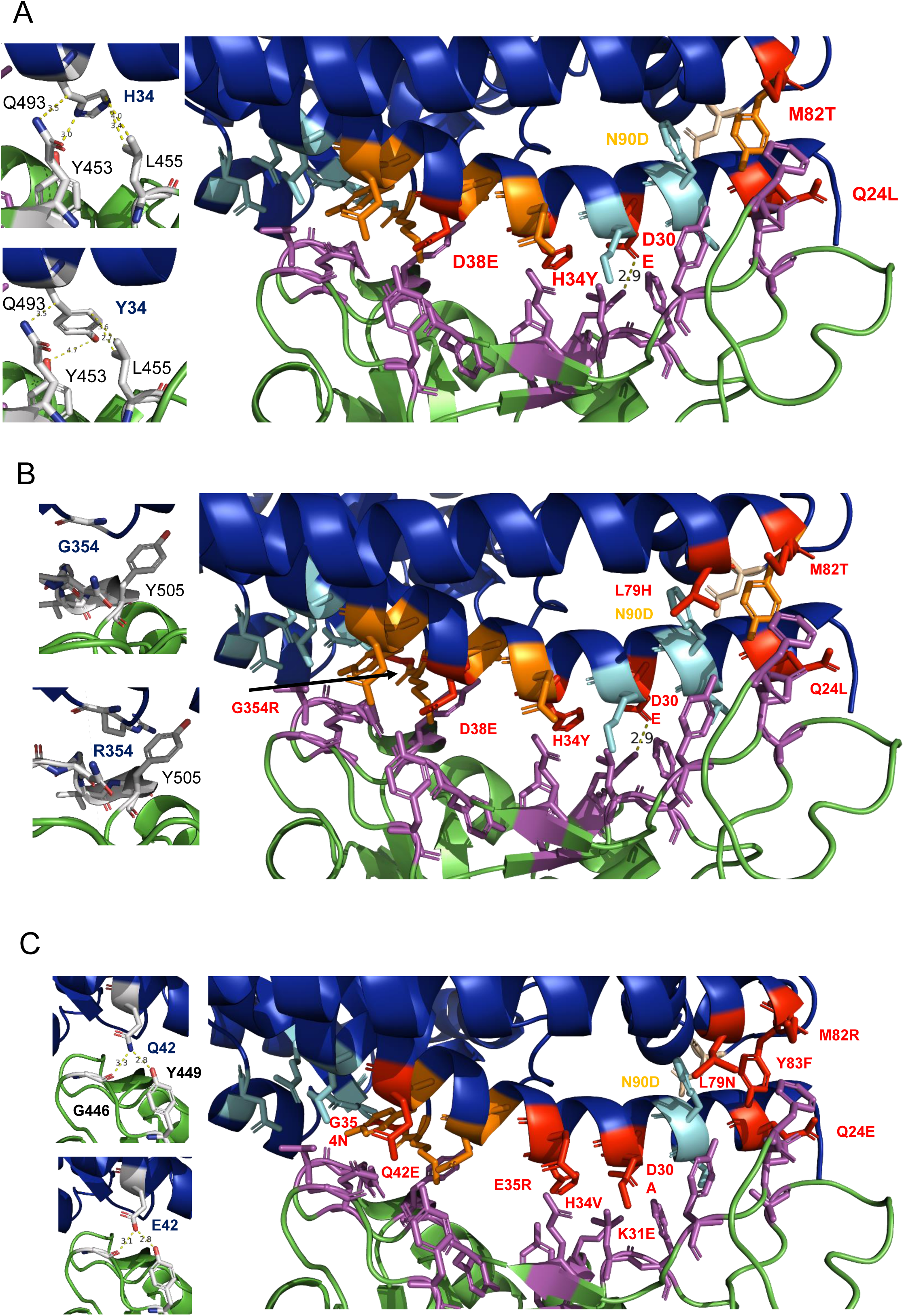
(A) Amino acid changes between human ACE2 and dog ACE2. (B) Amino acid changes between human ACE2 and ferret ACE2.

**Cat ACE2** contains only four changes (Gln24Leu, Asp30Glu, Asp38Glu, Met82Thr), which are also present in dog ACE2 (Table 1). For the first time, we analysed the ability of ACE2 proteins from pigs, dogs and cats binding to S and submitted this manuscript on 27^th^ of March. During the review process, results of infection experiments with animals in close contact with humans were posted at 31^th^ of March (https://www.biorxiv.org/content/10.1101/2020.03.30.015347v1.full.pdf), and it showed that SARS-CoV-2 replicates poorly in dogs, but efficiently in cats and was transmitted by droplets to naïve cats that ACE2 is linked to the fact (13). This indicated that S binding to ACE2 receptors is only the first step in the virus’s invasion, and ACE2 levels in different tissues may play an important role in viral transmission. Therefore, the risk of infection in animals requires continuous monitoring. The only residue in dogs, which is not changed in cat ACE2 is His34, which interacts with Tyr453, Leu455, and Gln493 in the centre of the interaction surface. It is exchanged by the slightly later Tyr residue in dog ACE2, which is still able to interact with the same residues in S (Fig. 3A, inset). The other difference is the loop of the N-glycosylation site at position 90.

There is also anecdotal evidence that **tigers** and **lions** in the Bronx Zoo of New York City were infected by SARS-CoV-2 (https://www.aphis.usda.gov/aphis/newsroom/news/sa_by_date/sa-2020/ny-zoo-covid-19). Therefore, we analysed the ACE2 gene from these wild cats. One amino acid difference was detected in ACE2 from cat and tiger, but the residues contacting S are identical explaining why tigers are also susceptible to SARS-CoV-2 infection. ACE2 from a lion has another conservative change relative to cat ACE2: His34 is substituted by Ser and it exhibits a loss of the N-glycosylation site at position 90, like dog ACE2.

### Comparison of the surface of human ACE2 and ACE2 from species suitable as animal models

Exploring potential therapies for COVID-19 animal models is urgently needed. Recent study showed that ferrets to be highly susceptible to infection with SARS-CoV-2 and even transmit virus to naïve contact animals, but also by droplets, albeit the latter route was less efficient (13, 30). **Ferret ACE2** exhibits the exact same five changes as dog ACE2, but also a substitution of Leu79 by His. In addition, a drastic change occurs at position 354, where the small glycine residue is replaced by a large and positively charged Arg residue (Fig. 3B). However, the Arg residue avoids clashing with large amino acids in S (Tyr505) by protruding backwards (Fig. 3B, inset).

The **Syrian hamster** (*Mesocricetus auratus*) is also shown to be susceptible to experimental infection and transmitted SARS-CoV-2 to close contact animals (31). Its ACE2 protein contains only two amino acid substitutions relative to human ACE2. His34 is exchanged to Gln and Met82 is replaced by Asn. Since residue 84 is also exchanged to a Ser, it creates a N-glycosylation site at position 82. A glycan attached to the same position prevents binding of S of SARS to rat ACE2.

**Guinea pig** (*Cavia porcellus*) might also serve as an animal model, but also common pets, especially of children. Guinea pig ACE2 contains seven amino acid changes at positions 24, 27, 31, 34, 38, 82, and 354 and thus more than ACE2 from ferrets or dogs. The glycosylation site at position 90 is preserved, but the site at position 322 is lost due to an Asn to Pro change (Table 1).

### Comparison of the interacting surface of human ACE2 and ACE2 from farm animals

Farm animals are also in close contact with humans and thus represent another risk group that might become infected by SARS-CoV-2. The ACE2 proteins from **chicken** contain ten amino acid changes compared with human ACE2 and lost the N-glycosylation site at position 90 (Fig. 3C). Some of the affected positions (Gln24, His34, Leu79, Met82, and Gly 353) are also exchanged in ACE2 proteins that serve as SARS-CoV-2 receptor, albeit often to different residues. Unique to all ACE2 proteins is the change of Gln24 by the negatively charged Glu residue, but the interaction with Gly446 and Tyr449 in S is probably preserved (Fig 3C, inset). Most of the other exchanges are likely to be more critical for binding to S. Tyr at position 83 is replaced by a Phe, which is not able to form a hydrogen bond with S. Two of the changes reverse the polarity of a charged amino acid: Lys31 is substituted by a negatively charged Glu and Glu35 is exchanged to a positively charged Arg. Finally, Asp30, which forms the salt bridge with S, is replaced by the uncharged residue Ala, which makes the ACE2 proteins from chicken and mice the only ones that are not able to form a salt bridge with S.

**Duck** ACE2 also contains ten amino acid substitutions, nine of them at the same position and eight to the same amino, including all the presumably important ones just discussed. Thus, it seems likely that the lack of susceptibility of chicken and ducks to experimental SARS-CoV-2 is due to the inability of the virus to bind to the avian ACE2 receptor (13).

The ACE2 protein of pigs can serve as SARS-CoV-2 receptor although it contains five amino acid changes in amino acids contacting the S protein (Fig. 2A). **Cattle** and **sheep** contain only two amino acid changes (Asp30Glu and Met82Thr) that are also present in pig ACE2 and even retain the N-glycosylation site at position 90 of human ACE2 that is lost in pig ACE2. It thus seems highly likely that ACE2 proteins of both species can function as SARS-CoV-2 receptor and experimental infection of these animals and surveillance is required to show whether they are susceptible to SARS-CoV-2. **Camel** is the animal reservoir of the Middle East Respiratory syndrome virus (MERS), which however uses dipeptidyl-peptidase 4 as protein receptor. We analysed the ACE2 gene from *Camelus bactrianus* and found, besides the two changes present in cattle and sheep, another substitution: the positively charged Lys31 located at the center of the binding site is exchanged by a negatively charged Glu residue, a change which is also present in guinea pig.

In summary, almost all mammalian species known to be susceptible to SARS-CoV-2 infection (cats and ferrets) have mutations in many amino acids in their ACE2 proteins. This suggests that these species, especially those in contact with humans, are at risk of contracting the virus and SARS-CoV-2 might establish itself in one of these animals thereby creating an additional animal reservoir. An exception is apparently the pig, which cannot be infected with SARS-CoV-2, although their ACE2 protein can function as SARS-CoV-2 receptor (12). In addition, dogs do not transmit the virus to naïve animals in close contact. We therefore investigated the level of ACE2 expression in different organs by q-RT-PCR. We found that pigs and dogs have the highest mRNA levels in kidney, but the level is also high in other internal organs, such as heart (pigs and dogs) and duodenum and liver (pigs) (Fig. 4). The relative mRNA levels in tissues relevant for SARS-CoV-2 infection, lung, trachea and turbinate are very low, 1,000-fold (dogs) or 10,000-fold (pigs) lower than in kidney. Among these organs, pigs exhibit the highest mRNA levels in trachea, around 20-fold less in lungs and very little in turbinates. In dogs the mRNA level is around 20-fold higher in lungs compared with the trachea, no mRNA was detected in turbinates. It is thus tempting to speculate that the low ACE2 levels in the respiratory tract prevent infection of pigs and the efficient replication and hence spread of the virus to contact animals in dogs. However, efficient receptor binding is only the first step in virus infection. Beyond that, numerous restriction factors have been identified in cells from various species that prevent successful virus replication. Our results mean more experiments and monitoring should be done to explore the source of animal infection.

**Fig. 4.**
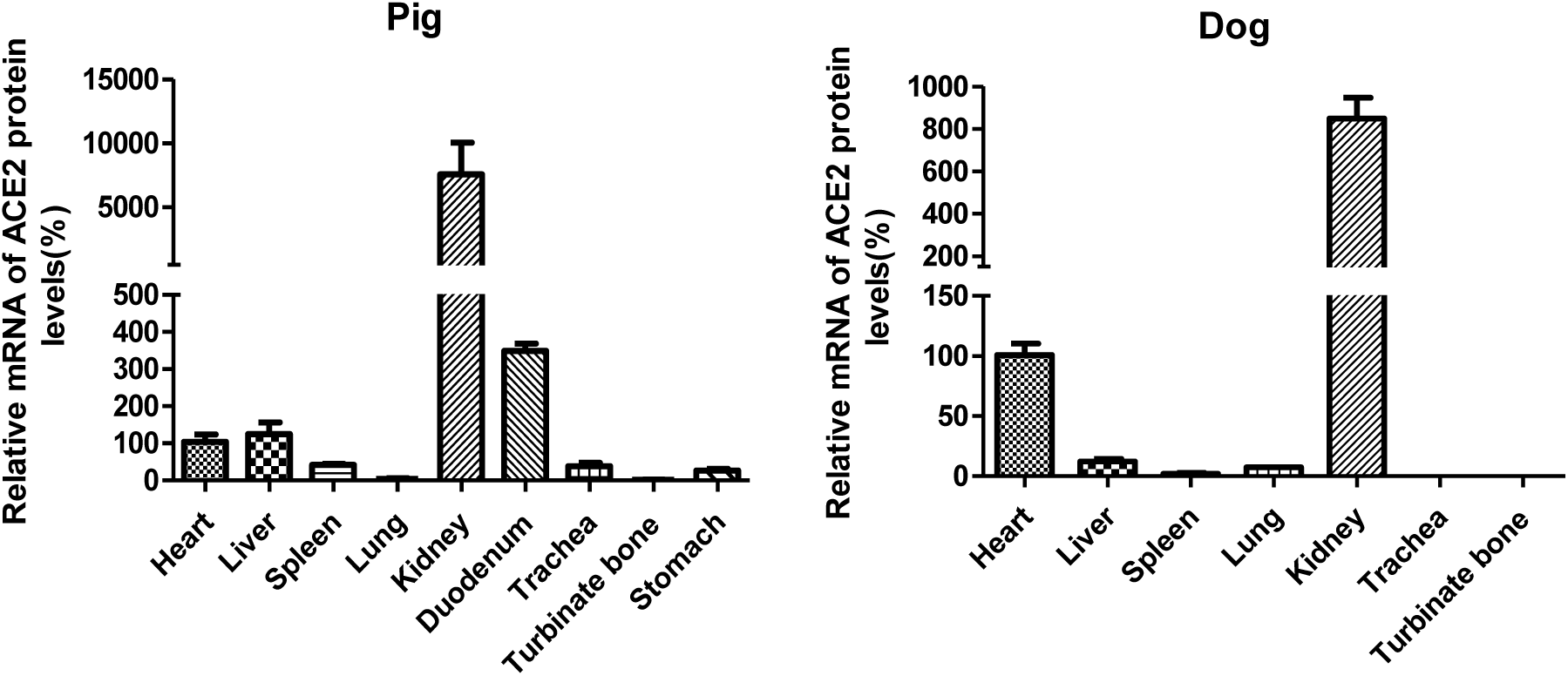
The relative expression of ACE2 in (A) pig and (B) dog tissues was determined by q-RT-PCR. (A) The relative expression of ACE2 in pig was determined in heart, liver, spleen, lung, kidney, duodenum, trachea, turbinate bone, and stomach. (B)The relative expression of ACE2 in dog was determined in heart, liver, spleen, lung, kidney, trachea, and turbinate bone. The experiments were repeated three times.

### Amino acids in ACE2 essential for binding of SARS-CoV-2

Based on an amino acid comparison of ACE2 proteins from animals that do not serve as SARS-CoV-2 receptor (mice) or are not susceptible to SARS-CoV-2 infection (chicken) with ACE2 proteins from animals which are infectable (cats, dogs, ferrets) or encode a ACE2 protein that confers susceptibility to SARS-CoV-2 infection (civet), the amino acids essential for binding of S can be deduced (Fig 5, Table 1). **Tyr83**, which forms hydrogen bonds with Asn487 and Tyr489 in S, is replaced by Phe in mice and duck. Phe has the same size and hydrophobicity but is not able to form hydrogen bonds with its side chain. All other animals exhibit at position 83 a Tyr residue. Note, however, that Tyr489 might be shielded by the acquisition of an N-glycosylation site at position 82 as it occurs in ACE2 of mice and in some bat sequences. The other residues are located in the center of the interaction surface. **Lys31**, which forms van-der-Waals contacts with Y489 and F456 is exchanged by a neutral Asn reside in mice and even to a negatively charged Glu in chicken. **Glu35** forms hydrogen bonds with Gln493 in S. It is replaced by a positively charged Arg in chicken, ducks, and some bat sequences. **Lys353**, which forms hydrogen bonds with its side chain to Tyr495, Gly496, and Gly502, is replaced by the smaller His residue in mice, which presumably cannot form these hydrogen bonds. ACE2 of all other animals (including chicken) contain a Lys at this position. In support of this hypothesis, a single Lys353Ala mutation was shown to abolish the ACE-S interaction (32). Finally, also important is the centrally located **Asp30,** which is substituted by Asn in mice and chicken. Asn has the same size as Asp, but is uncharged and thus unable to sustain the salt bridge with Lys417 in S. ACE2 proteins from all other animals either retain Asp or it is substituted by the negatively charged, but slightly larger Glu residue, which is probably also able to from the salt bridge.

**Fig. 5.**
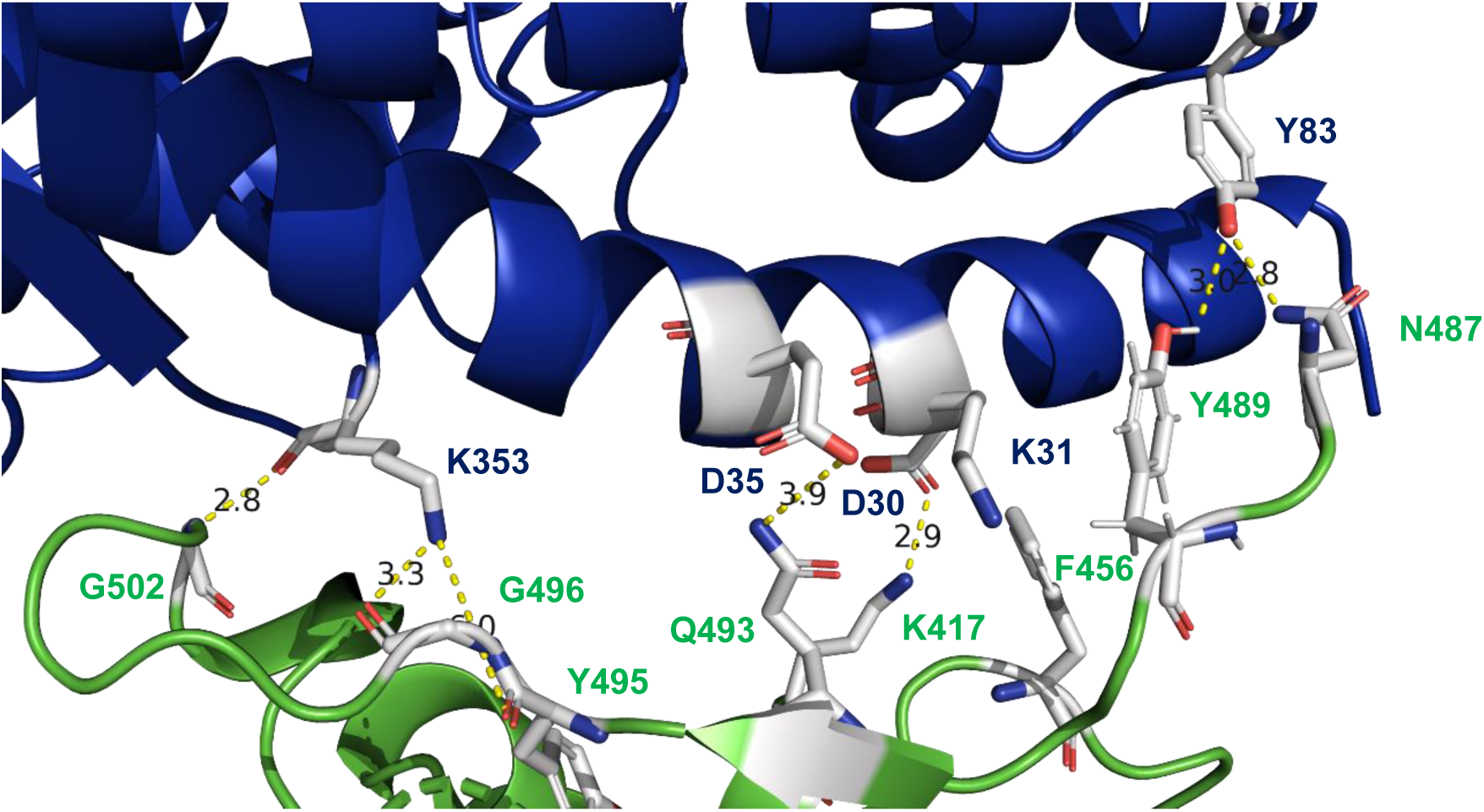
Amino acids in ACE2 important for binding to S of SARS-CoV-2. Y83 forms hydrogen bonds to N487 and Y489, which are probably weakened by the conservative change to F. K353 forms hydrogen bonds to G502, G496, and Y495, the latter two are probably destroyed by the change to His and the interacting amino acid changes between human and mouse ACE2. Amino acid changes in human ACE2 compared with mouse ACE2 are highlighted in red. P84 in human ACE does not interact with amino acids in S of SARS-CoV-2 but might affect the secondary structure. N90T and N 322H destroy the N-glycosylation site in human ACE2.

Note that positions 330, 355, and 357 are conserved through all ACE2 proteins we analyzed here and thus their relevance for binding to S cannot be estimated (Table 1). In addition, binding of S to ACE2 might not follow a simple “lock-and-key” principle. The mouse-adapted SARS-CoV strain MA15 contains a single amino acid exchange in the S protein relative to the Urbani strain; Tyr at position 346 is replaced by a His residue (33). Tyr 346 forms hydrogen bonds with Asp38 and Gln42 in human ACE2 (Fig. 1B), but mouse ACE2 also contains Asp and Gln at positions 38 and 42, respectively. His is also able to serve as hydrogen-bond donor or acceptor, but its side chain is shorter, and it is not obvious how this exchange enhances binding to mouse ACE2.

### Comparison of amino acids in the RBM of SARS-CoV-2 with bat and pangolin coronavirus

It has been reported that SARS-CoV-2 derived from a bat virus, but parts of the S protein exhibit a higher nucleotide similarity to a virus from pangolin. Therefore, we aligned the whole amino acid sequence of S from SARS-CoV-2 (first Wuhan isolate) with sequences from the bat and the pangolin virus (Supplementary Fig. 1). As noted before, S from SARS-CoV-2 contains an insertion of four amino acids (PRRA) that creates a polybasic cleavage site recognized by the ubiquitous protease furin (20, 32). Insertion of amino acids at the S1/S2 junction can occur also in bats, since the novel bat-derived virus RmYN02 contains the insertion of amino acid PAA, which does not create a polybasic cleavage site (34). The following S2 subunit is almost completely conserved among all three viruses, it exhibits nine mostly conservative amino acid substitutions in S of the pangolin virus, and only two in S of the bat virus. The N-terminus of the S protein until residue 400 is also highly similar between S of SARS-CoV-2 and the bat virus. There are only six amino acid exchanges in the bat virus, two of them affecting N-glycosylation sites, whereas S from the pangolin virus contains 101 amino acid differences compared with S of SARS-CoV-2 (Supplementary Fig. 1). However, from residue 401 to 518, which contains the receptor binding domain of S, the homology reverses (Table 2).

**Table 2.**
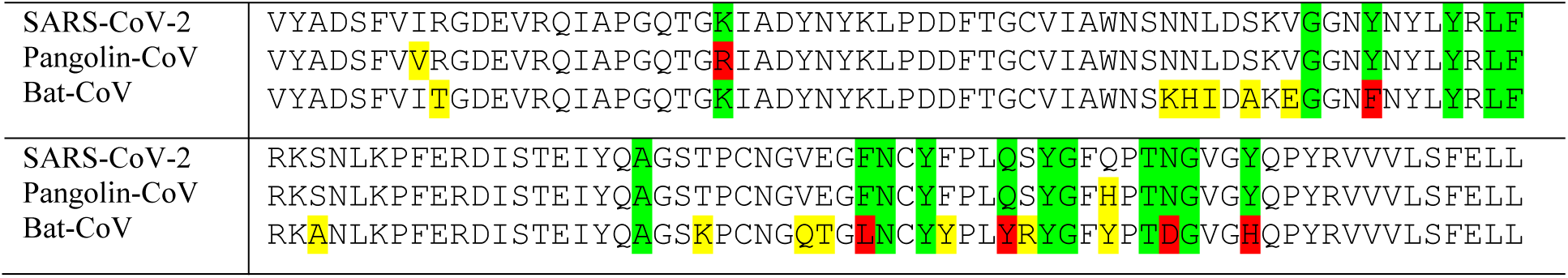
Amino acid alignment of S receptor binding region

The bat virus contains 18 amino acid substitutions, five of them involve amino acids that contact human ACE2 (white sticks in Fig. 6A). Located at the periphery of the interaction surface is Phe486, which interacts with Leu79 in ACE2. It is replaced by a Leu, that is also a large and hydrophobic residue. At the other side of the interaction surface located is Asn501, which forms a hydrogen bond with Tyr41 in ACE2 and is replaced by the negatively charged Glu residue. The other three exchanged amino acids are located in the centre of the ACE2 binding site. Gln493, which forms a hydrogen bond with Glu35, is replaced by a Tyr residue. Tyr449, which forms a hydrogen bond with Gln42, is replaced by Phe, which has the same size, but cannot form (or only weak) hydrogen bonds. The most drastic exchange is probably Tyr505, which forms a hydrogen bond with Gln42 and is exchanged by a much smaller His residue.

**Fig. 6.**
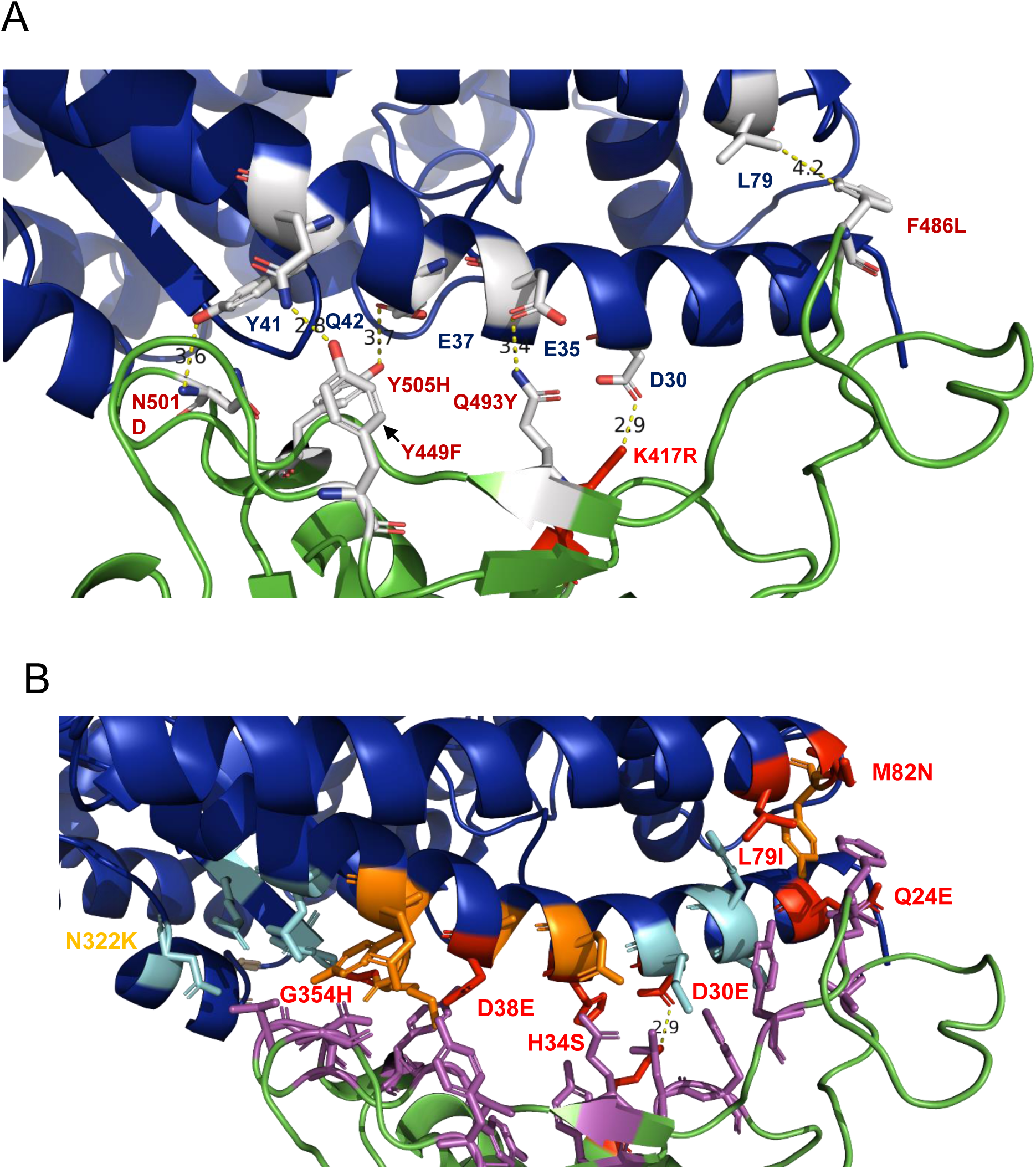
(A) Amino acid changes in the RBD of S from SARS-CoV-2 and viruses from bats (white sticks) and pangolin (red stick). Hydrogen bonds between S from SARS-CoV-2 and the corresponding amino acids in ACE2, which are likely to be weakened or destroyed by the changes, are shown as dotted lines. (B) Amino acid changes between human ACE2 and pangolin ACE2.

In contrast, S from the pangolin virus contains only three exchanges relative to SARS-CoV-2, and only one of them involves an amino acid that contacts ACE2, the substitution of Lys 417 by an Arg (red stick in Fig. 6A). Since both are basic amino acids, the important salt bridge is probably preserved. In summary, the RBD from the pangolin coronavirus is much better suited to bind to human ACE2 compared to the bat virus. From this point of view, it seems possible that SARS-CoV-2 might have acquired the RBD from a pangolin coronavirus to achieve bat-to-human transmission.

### Comparison of the interacting surface of ACE2 proteins of putative intermediate hosts

We therefore asked whether ACE2 of **pangolin** (*Manis javanica*) contains amino acids in its interaction surface that might closely resemble those of humans. In that case, a precursor of SARS-CoV-2 acquired a RBD from a pangolin virus by recombination which is then already adapted to replicate both in pangolins and in humans. However, pangolin ACE2 contains seven changes compared with human ACE2 (Fig. 6B). In addition, the N-glycosylation site at position 322 of human ACE2 is lost due to an change of Asn to Lys. Some of the changes, Asp30Glu, His34Ser, Asp38Glu, and Leu79Ile, occur also in ACE2 proteins from animals that can interact with S. The three other variable positions are also exchanged in other animals, but mostly to other residues. Gly354 is replaced by a small His residue and not by the larger Arg, that is present in ferret ACE2. Gln24, which interacts with Asn 487 in S, is replaced by a negatively charged Glu residue. Finally, Met82, that interacts with Phe486 in S, is replaced by the larger Asn residue. Although none of the amino acid changes might prevent binding to S, it nevertheless appears that pangolin ACE2 is not especially equipped to serve as receptor for SARS-CoV-2.

Other potential intermediate hosts are civets and **raccoon dogs** (*Nyctereutes procyonoides*). ACE2 of civets has been shown to confer susceptibility to SARS-CoV-2 infection in cell culture, although it contains seven changes relative to human ACE2 in the amino acids contacting S (Fig. 2B). The ACE2 protein from racoons contains only five substitutions, Gln24Leu, Asp30Glu, His34Tyr, Asp38Glu, and Met82Thr, which are also present in civet ACE2. It is also identical in these residues to ACE2 from dogs, which are susceptible to SARS-CoV-2 infection but do not spread the virus.

In summary, none of ACE2 proteins of any of the discussed intermediate hosts seems to be especially equipped to attach to S of SARS-CoV-2, but the least number of changes occur in ACE2 of racoon dogs.

### Glycosylation sites and a putative integrin binding motif in S proteins from SARS-CoV-2 and bat and pangolin virus

It has previously been shown that a RGD motif (403–405: Arg-Gly-Asp) is present in the receptor-binding domain of the spike proteins of all SARS-CoV-2 sequences (35). This sequence mediates attachment of several viruses to integrins, which thus might serve as an additional receptor for SARS-CoV-2. Two RGD motifs are present in S of pangolin CoV, one at the same site, another at amino acids 246-249, but S of bat coronavirus (RaTG13) contains no RGD motif (Fig. 7A).

**Fig. 7.**
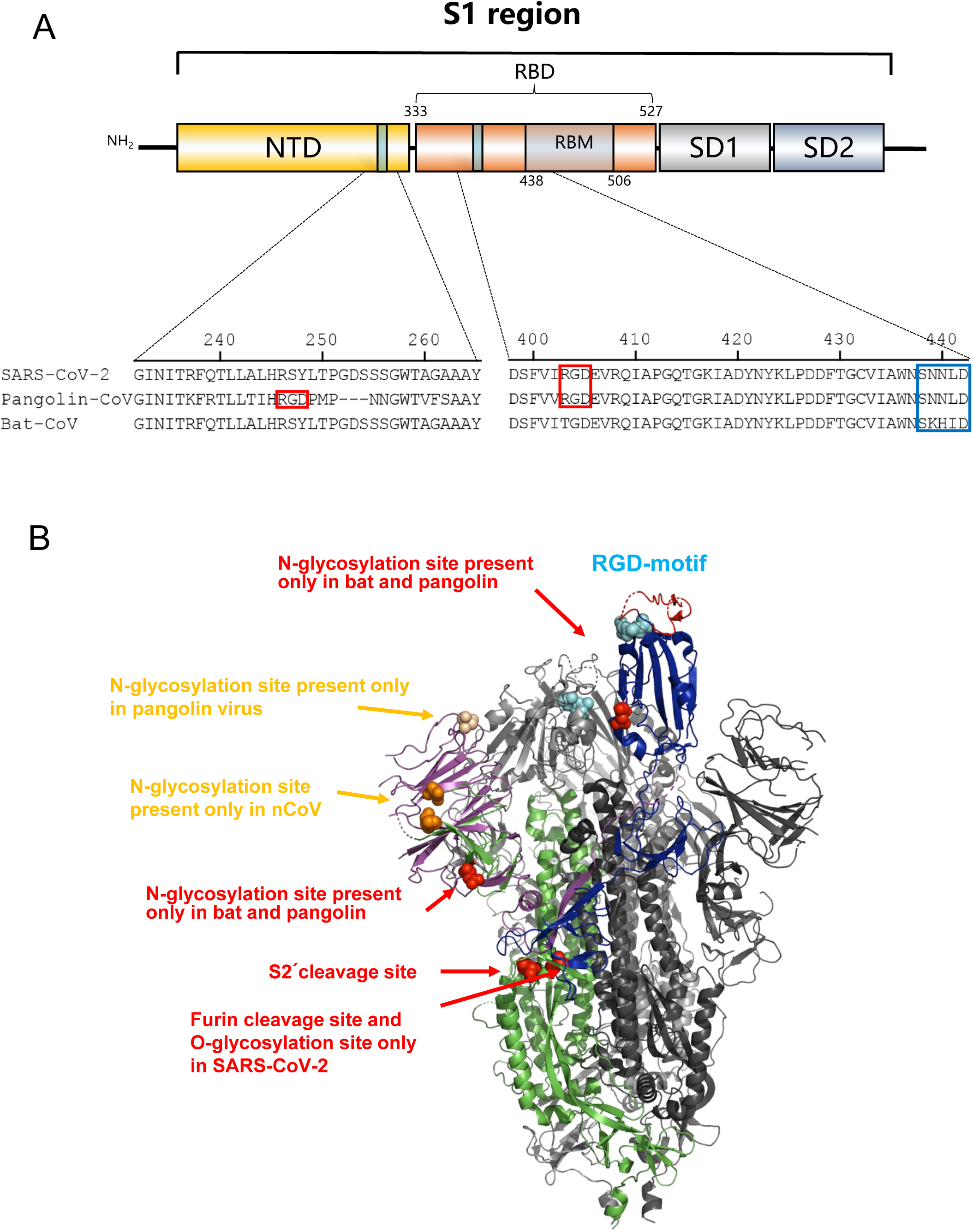
Distribution of glycosylation sites and RGD motifs in the S from SARS-CoV-2, bat and pangolin CoV. (A) Schematic representation of SARS-CoV-2 S1 protein. The sequences of SARS-CoV-2, bat and pangolin coronavirus were aligned using MegAlign. The RGD motif is highlighted in red. The RBM starting position is highlighted in blue. (B) Cryo-EM structure of S from SARS-CoV-2 (14). N-glycosylation sites unique for SARS-CoV-2, bat and/or pangolin virus are highlighted as red or orange spheres. The amino acid sequences RGD present in S of SARS-CoV-2 and the pangolin virus is labeled as cyan sphere. In a monomer in the open conformation it is located near the ACE2 binding sites on top of the molecule. The location of the RGD motif in a monomer in a closed conformation (grey) is also indicated. The location of the furin cleavage site present only in S of SARS-CoV-2 and of the S2’cleavage site is also highlighted as red spheres.

Bulky carbohydrates attached to the S protein might mask antibody epitopes, interfere with receptor binding and/or with proteolytical cleavage that is required to prime the protein to execute its membrane fusion activity. S of SARS-CoV-2 contains 22 N-glycosylation sites (NXS/T), nine in S2 and 13 in the S1 subunit, which are all glycosylated with almost 100% stoichiometry if the ectodomain of S is expressed in mammalian cells (36).

A total of 21 glycosylation sites are conserved among all three viruses. S of SARS-CoV-2 contains a unique glycosylation site in the N-terminal domain at position 74 in a loop of the N-terminal S1 domain (Fig. 7B). This region (aa 60-80) is dissimilar to the pangolin virus, but identical to S of the bat virus, except the glycosylation site (NGI). Two sites are present only in S of the bat and pangolin coronavirus, but not in SARS-CoV-2. One is also located in the N-terminal domain at position 30, in a region (aa 15-49) which is identical to S of SARS-CoV-2 virus, except the glycosylation sequon, which is NSS in S of bat, but NSF in S of SARS-CoV-2. The N-terminal domain has a galectin-like folding and is known in other coronaviruses to bind to carbohydrates on the cell surface, i. e. using them as an attachment factor as the first step of virus entry (17). Most interestingly is a site at position 370 which localizes near the ACE2 receptor binding domain. This residue is located in a region (aa 275-400) of high amino acid identity among all three viruses. Thus, it is tempting to speculate that this site was lost during adaption of SARS-CoV-2 to humans, in order to get better access to the ACE2 receptor.

As suggested before, the insertion of amino acids at the cleavage site between the S1/S2 subunits of SARS-CoV 2 creates three potential GalNAc O-glycosylation sites, which are not predicted for S of the bat or pangolin virus (37). Attachment of GalNac to serine and threonine residues is catalysed by up to 20 different Gal-Nac transferases having different, partially overlapping substrate specificities. The sugar residues are then elongated by other glycosyltransferases thereby creating long carbohydrate chains. Interestingly, whether a certain O-glycoslyation site is used is cell-type dependent and O-glycans attached near furin cleavage site have been shown to affect processing (38, 39). It is thus tempting to speculate that the usage of an O-glycosylation site might determine whether S is cleaved in a certain cell. Since processing by furin is essential for entry of SARS-CoV-2 into cells that lack cathepsin proteases, O-glycosylation might affect cell tropism and hence virulence of SARS-CoV-2 (40). This is somewhat reminiscent of hemagglutinin (HA) of an avian Influenza A virus where the loss of a N-glycosylation sequon near a polybasic cleavage site allowed processing of HA thereby generating a highly virulent strain (41).

## MATERIALS AND METHODS

### Sequence analysis

The S protein of SARS-CoV-2 and bat coronavirus RaTG13 were obtained from GenBank (accession numbers QHD43416.1 and QHR63300.2, respectively). The pangolin coronavirus nucleotide sequence was obtained from GISAID (EPI_ISL_410721) and translated into protein. Sequences were aligned using MEGA7.0. A total of 23 ACE2 protein sequences of 20 mammal including human, dog, cat, guinea pig, hamster, mouse, pig, rabbit, cattle. sheep, ferret, raccoon dog, bat (*Rhinolophus sinicus*), civet, pangolin, tiger, lion, camel, chicken and duck were download from GenBank. The ACE2 protein sequences of dog (MT269670) and cat (MT269670) were obtained from this study (Table 1). All the sequences were aligned using MEGA7.0.

### Structural analysis

The software PyMol was used to create the figures of the Cryo-EM structure of S from SARS-CoV-2 (pdb file 6VSB) and from the crystal structure of its receptor-binding domain bound to human ACE2 (pdb file 6M0J). The amino acids which are mentioned in this publication to mediate contact between S and ACE2 are shown as sticks, whereas the remainder of the molecules are shown as cartoons. The integrated measuring wizard was used to determine the distance between two atoms, which is shown as dotted lines also indicating the distance in Angström. Exchange of certain amino acids was performed with the mutagenesis tool. Among the different possible rotamers of the mutated amino acid side chain the one was chosen that exhibits no clashes with neighboring amino acids.

### Sample collection and nucleic acid extraction

Sample collection and processing were done at the Institute of Military Veterinary Medicine and Nanjing Agricultural University for animal welfare. All the experiments were approved by the committee on the ethics of animal experiments. Samples of dog and pig including heart, liver, spleen, lung, kidney, duodenum, bronchus and turbinate, were used for quantitative real-time PCR (q-RT-PCR) to determine the content of mRNA encoding ACE2. The kidney of a beagle or *Felis catus* were used for PCR amplification of the full length of ACE2. A total of 50 mg of each sample was ground into powder in liquid nitrogen and transferred to DEPC treated EP tube before liquid nitrogen volatilizes. Total RNA was extracted using 1 mL TRIzol (Nanjing Vazyme biotechnology), and 1 μg cDNA was synthesized using the RevertAid First Strand cDNA Synthesis Kit (Thermo Fisher Scientific) according to the manufacturer’s instructions.

### Amplification and detection of ACE2 by q-RT-PCR

The primers used to amplify the full length of ACE2 gene sequence of cat and dog by PCR included, Cat-ACE2-F: 5 ′ -AAGAGCTCATGTCAGGCTCTTTCTGGCTC-3 ′, Cat-ACE2-R: 5 ′ - TTGGTACCCTAAAATGAAGTCTGAACATCATCA-3 ′ and Dog-ACE2-F: 5 ′ - AAGAGCTCATGTCAGGCTCTTCCTGGCT-3 ′, Dog-ACE2-R: 5 ′ - TTGGTACCCTAAAATGAAGTCTGAGCATCATC-3′. The 25 μL PCR reaction mixtures contained, 1 μL each primer, 1 μL cDNA template, 12.5 μL 2 × Phanta Max Buffer, 0.5 μL dNTP Mix (10 mM each), 0.5 μL phanta Max Super-Fidelity DNA Polymerase (Nanjing Vazyme biotechnology Co., Ltd), and 8.5 μL ddH_2_O. PCR conditions were as follows: pre-denaturation at 95 °C for 5 min, 35 cycles of 95 °C for 30 s, 62 °C for 30 s, 72 °C for 2 min and 30 s, and a final extension at 72 °C for 10 min. Positive amplicons were sequenced by Comate Biosciences Company Limited (Changchun, Jilin) using an ABI 3730 sequencer, and consensus sequences were obtained from at least three independent determinations.

Q-RT-PCR was used to determine the content of ACE2 in different tissues of dog and pig. The total volume of q-RT-PCR reaction was 20 μL, including 10 μL 2x Aceq qPCR mixture (Nanjing Vazyme biotechnology), 0.4 μL of forward and reverse primer, 2 μL of template, and the remaining volume of nuclease free water. The following procedure was used for amplification on a Roche lightcycle® 96 instrument: 95 °C 600 s; 45 cycles of 95 °C 10 s, 55 °C 10 s, 72 °C 20 s. β-Actin was selected as the internal parameter. The primers for ACE2 genes of pig and dog were: Sus-ACE2-F 5′-TTGATGGAAGATGTGGAGCG-3′, Sus-ACE2-R 5′-CCACATATCGCCAAGCAAATG-3′, Sus-β-actin-F 5′-CTCCATCATGAAGTGCGACGT-3′, Sus-β-actin-R 5′-GTGATCTCCTTCTGCATCCTGTC-3′, Dog-ACE2-F 5′-GGTGGATGGTCTTTAAGGGTG-3′, Dog-ACE2-R 5′-ACATGGAACAGAGATGCAGG-3′, Dog-β-actin-F 5′-GGACTTCGAGCAGGAGATGG-3′, Dog-β-actin-R 5′-TTCCATGCCCAGGAAGGAAG-3′.

### Statistical analysis

Data is expressed as mean values ± s.d. or s.e.m. Statistical analyses were performed using Student’s t-test using GraphPad Prism 5 software.

## ACKNOWLEDGMENTS

This work was financially supported by the National Key Research and Development Program of China (2017YFD0500101), the Fundamental Research Funds for the Central Universities Y0201900459, the Young Top-Notch Talents of National Ten-Thousand Talents Program, Sino German cooperation and exchange project in international cooperation and Cultivation Project in 2019. Experimental work in the lab of M.V. is financed by the German Research Foundation (DFG).

