## Supplementary Fig. 1 for "Comparison of SARS-CoV-2 spike protein binding to human, pet, farm animals, and putative intermediate hosts ACE2 and ACE2 receptors"

```
Pangolin-CoV  MLFFFFLHFALVNSQCVNLTGRAAIQPSFTNSSQRGVYYPDTIFRSNTLVLSQGYFLPFY	60
SARS-CoV-2    -MFVFLVLLPLVSSQCVNLTTRTQLPPAYTNSFTRGVYYPDKVFRSSVLHSTQDLFLPFF	59
Bat-CoV       -MFVFLVLLPLVSSQCVNLTTRTQLPPAYTNSSTRGVYYPDKVFRSSVLHLTQDLFLPFF	59
               :*.*:: : **.******* *: : *::***  *******.:***..*  :*. ****:

Pangolin-CoV  SNVSWYYALTKTN-SAEKRVDNPVLDFKDGIYFAATEKSNIVRGWIFGTTLDNTSQSLLI	119
SARS-CoV-2    SNVTWFHAIHVSGTNGTKRFDNPVLPFNDGVYFASTEKSNIIRGWIFGTTLDSKTQSLLI	119
Bat-CoV       SNVTWFHAIHVSGTNGIKRFDNPVLPFNDGVYFASTEKSNIIRGWIFGTTLDSKTQSLLI	119
              ***:*::*:  :. .. **.***** *:**:***:******:**********..:*****

Pangolin-CoV  VNNATNVIIKVCNFQFCYDPYLSGYYHN-NKTWSTREFAVYSSYANCTFEYVSKSFMLDI	178
SARS-CoV-2    VNNATNVVIKVCEFQFCNDPFLGVYYHKNNKSWMESEFRVYSSANNCTFEYVSQPFLMDL	179
Bat-CoV       VNNATNVVIKVCEFQFCNDPFLGVYYHKNNKSWMESEFRVYSSANNCTFEYVSQPFLMDL	179
              *******:****:**** **:*. ***: **:*   ** ****  ********: *::*:

Pangolin-CoV  AGKSGLFDTLREFVFRNVDGYFKIYSKYTPVNVNSNLPIGFSALEPLVEIPAGINITKFR	238
SARS-CoV-2    EGKQGNFKNLREFVFKNIDGYFKIYSKHTPINLVRDLPQGFSALEPLVDLPIGINITRFQ	239
Bat-CoV       EGKQGNFKNLREFVFKNIDGYFKIYSKHTPINLVRDLPPGFSALEPLVDLPIGINITRFQ	239
               **.* *..******:*:*********:**:*:  :** *********::* *****:*:

Pangolin-CoV  TLLTIHRGDPMP---NNGWTVFSAAYYVGYLAPRTFMLNYNENGTITDAVDCALDPLSEA	295
SARS-CoV-2    TLLALHRSYLTPGDSSSGWTAGAAAYYVGYLQPRTFLLKYNENGTITDAVDCALDPLSET	299
Bat-CoV       TLLALHRSYLTPGDSSSGWTAGAAAYYVGYLQPRTFLLKYNENGTITDAVDCALDPLSET	299
              ***::**.   *   ..***. :******** ****:*:********************:

Pangolin-CoV  KCTLKSLTVEKGIYQTSNFRVQPTESIVRFPNITNLCPFGEVFNATTFASVYAWNRKRIS	355
SARS-CoV-2    KCTLKSFTVEKGIYQTSNFRVQPTESIVRFPNITNLCPFGEVFNATRFASVYAWNRKRIS	359
Bat-CoV       KCTLKSFTVEKGIYQTSNFRVQPTDSIVRFPNITNLCPFGEVFNATTFASVYAWNRKRIS	359
              ******:*****************:********************* *************

Pangolin-CoV  NCVADYSVLYNSTSFSTFKCYGVSPTKLNDLCFTNVYADSFVVRGDEVRQIAPGQTGRIA	415
SARS-CoV-2    NCVADYSVLYNSASFSTFKCYGVSPTKLNDLCFTNVYADSFVIRGDEVRQIAPGQTGKIA	419
Bat-CoV       NCVADYSVLYNSTSFSTFKCYGVSPTKLNDLCFTNVYADSFVITGDEVRQIAPGQTGKIA	419
              ************:*****************************: *************:**

Pangolin-CoV  DYNYKLPDDFTGCVIAWNSNNLDSKVGGNYNYLYRLFRKSNLKPFERDISTEIYQAGSTP	475
SARS-CoV-2    DYNYKLPDDFTGCVIAWNSNNLDSKVGGNYNYLYRLFRKSNLKPFERDISTEIYQAGSTP	479
Bat-CoV       DYNYKLPDDFTGCVIAWNSKHIDAKEGGNFNYLYRLFRKANLKPFERDISTEIYQAGSKP	479
              *******************:::*:* ***:*********:******************.*

Pangolin-CoV  CNGVEGFNCYFPLQSYGFHPTNGVGYQPYRVVVLSFELLNAPATVCGPKQSTNLVKNKCV	535
SARS-CoV-2    CNGVEGFNCYFPLQSYGFQPTNGVGYQPYRVVVLSFELLHAPATVCGPKKSTNLVKNKCV	539
Bat-CoV       CNGQTGLNCYYPLYRYGFYPTDGVGHQPYRVVVLSFELLNAPATVCGPKKSTNLVKNKCV	539
              ***  *:***:**  *** **:***:*************:*********:**********
Pangolin-CoV  NFNFNGLTGTGVLTESSKKFLPFQQFGRDIADTTDAVRDPQTLEILDITPCSFGGVSVIT	595
SARS-CoV-2    NFNFNGLTGTGVLTESNKKFLPFQQFGRDIADTTDAVRDPQTLEILDITPCSFGGVSVIT	599
Bat-CoV       NFNFNGLTGTGVLTESNKKFLPFQQFGRDIADTTDAVRDPQTLEILDITPCSFGGVSVIT	599
              ****************.*******************************************

Pangolin-CoV  PGTNTSNQVAVLYQDVNCTEVPVAIHADQLTPTWSVYSTGSNVFQTRAGCLIGAEHVNNS	655
SARS-CoV-2    PGTNTSNQVAVLYQDVNCTEVPVAIHADQLTPTWRVYSTGSNVFQTRAGCLIGAEHVNNS	659
Bat-CoV       PGTNASNQVAVLYQDVNCTEVPVAIHADQLTPTWRVYSTGSNVFQTRAGCLIGAEHVNNS	659
              ****:***************************** *************************

Pangolin-CoV  YECDIPIGAGICASYQTQTNS----RSVSSQAIIAYTMSLGAENSVAYANNSIAIPTNFT	711
SARS-CoV-2    YECDIPIGAGICASYQTQTNSPRRARSVASQSIIAYTMSLGAENSVAYSNNSIAIPTNFT	719
Bat-CoV       YECDIPIGAGICASYQTQTNS----RSVASQSIIAYTMSLGAENSVAYSNNSIAIPTNFT	715
              *********************    ***:**:****************:***********

Pangolin-CoV  ISVTTEILPVSMTKTSVDCTMYICGDSIECSNLLLQYGSFCTQLNRALTGIAVEQDKNTQ	771
SARS-CoV-2    ISVTTEILPVSMTKTSVDCTMYICGDSTECSNLLLQYGSFCTQLNRALTGIAVEQDKNTQ	779
Bat-CoV       ISVTTEILPVSMTKTSVDCTMYICGDSTECSNLLLQYGSFCTQLNRALTGIAVEQDKNTQ	775
              *************************** ********************************

Pangolin-CoV  EVFAQVKQIYKTPPIKDFGGFNFSQILPDPSKPSKRSFIEDLLFNKVTLADAGFIKQYGD	831
SARS-CoV-2    EVFAQVKQIYKTPPIKDFGGFNFSQILPDPSKPSKRSFIEDLLFNKVTLADAGFIKQYGD	839
Bat-CoV       EVFAQVKQIYKTPPIKDFGGFNFSQILPDPSKPSKRSFIEDLLFNKVTLADAGFIKQYGD	835
              ************************************************************

Pangolin-CoV  CLGDIAARDLICAQKFNGLTVLPPLLTDEMIAQYTSALLAGTITSGWTFGAGAALQIPFA	891
SARS-CoV-2    CLGDIAARDLICAQKFNGLTVLPPLLTDEMIAQYTSALLAGTITSGWTFGAGAALQIPFA	899
Bat-CoV       CLGDIAARDLICAQKFNGLTVLPPLLTDEMIAQYTSALLAGTITSGWTFGAGAALQIPFA	895
              ************************************************************

Pangolin-CoV  MQMAYRFNGIGVTQNVLYENQKLIANQFNSAIGKIQDSLSSTASALGKLQDVVNQNAQAL	951
SARS-CoV-2    MQMAYRFNGIGVTQNVLYENQKLIANQFNSAIGKIQDSLSSTASALGKLQDVVNQNAQAL	959
Bat-CoV       MQMAYRFNGIGVTQNVLYENQKLIANQFNSAIGKIQDSLSSTASALGKLQDVVNQNAQAL	955
              ************************************************************

Pangolin-CoV  NTLVKQLSSNFGAISSVLNDILSRLDKVEAEVQIDRLITGRLQSLQTYVTQQLIRAAEIR	1011
SARS-CoV-2    NTLVKQLSSNFGAISSVLNDILSRLDKVEAEVQIDRLITGRLQSLQTYVTQQLIRAAEIR	1019
Bat-CoV       NTLVKQLSSNFGAISSVLNDILSRLDKVEAEVQIDRLITGRLQSLQTYVTQQLIRAAEIR	1015
              ************************************************************

Pangolin-CoV  ASANLAATKMSECVLGQSKRVDFCGKGYHLMSFPQSAPHGVVFLHVTYVPSQEKNFTTTP	1071
SARS-CoV-2    ASANLAATKMSECVLGQSKRVDFCGKGYHLMSFPQSAPHGVVFLHVTYVPAQEKNFTTAP	1079
Bat-CoV       ASANLAATKMSECVLGQSKRVDFCGKGYHLMSFPQSAPHGVVFLHVTYVPAQEKNFTTAP	1075
              **************************************************:*******:*

Pangolin-CoV  AICHEGKAHFPREGVFVSNGTHWFVTQRNFYEPQIITTDNTFVSGSCDVVIGIVNNTVYD	1131
SARS-CoV-2    AICHDGKAHFPREGVFVSNGTHWFVTQRNFYEPQIITTDNTFVSGNCDVVIGIVNNTVYD	1139
Bat-CoV       AICHDGKAHFPREGVFVSNGTHWFVTQRNFYEPQIITTDNTFVSGSCDVVIGIVNNTVYD	1135
              ****:****************************************.**************

Pangolin-CoV  PLQPELDSFKEELDKYFKNHTSPDVDLGDISGINASVVNIQKEIDRLNEVAKNLNESLID	1191
SARS-CoV-2    PLQPELDSFKEELDKYFKNHTSPDVDLGDISGINASVVNIQKEIDRLNEVAKNLNESLID	1199
Bat-CoV       PLQPELDSFKEELDKYFKNHTSPDVDLGDISGINASVVNIQKEIDRLNEVAKNLNESLID	1195
              ************************************************************

Pangolin-CoV  LQELGKYEQYIKWPWYIWLGFIAGLIAIIMVTIMLCCMTSCCSCLKGCCSCGSCCKFDED	1251
SARS-CoV-2    LQELGKYEQYIKWPWYIWLGFIAGLIAIVMVTIMLCCMTSCCSCLKGCCSCGSCCKFDED	1259
Bat-CoV       LQELGKYEQYIKWPWYIWLGFIAGLIAIIMVTIMLCCMTSCCSCLKGCCSCGSCCKFDED	1255
              ****************************:*******************************

Pangolin-CoV  DSEPVLKGVKLHYT	1265
SARS-CoV-2    DSEPVLKGVKLHYT	1273
Bat-CoV       DSEPVLKGVKLHYT	1269
              **************
```

**Supplementary Fig. 1** Sequence alignment of S proteins from SARS-CoV-2 (Wuhan strain, GenBank: QHD43416.1), Bat coronavirus (Bat-CoV) RaTG13 (GenBank: QHR63300.2) and Pangolin coronavirus (Pangolin-CoV) (EPI_ISL_410721). The Pangolin coronavirus nucleotide sequence was obtained from GISAID, and translated into protein. The N-terminal cleaved signal peptide and the C-terminal transmembrane region are underlined. Potential N-glycosylation sites (NXS/T) are highlighted in yellow. Amino acid exchanges relative to S of SARS-CoV-2 are highlighted in blue. The insertion of the amino acids PRRA in S of SARS-CoV-2 at the boundary between S1 and S2 is highlighted in red and the RGD motif is highlighted in purple. Putative O-glycosylation sites at the S1/S2 junction of SARS-CoV-2 as predicted by the NetOGlyc 4.0 Server (<http://www.cbs.dtu.dk/services/NetOGlyc/>) are highlighted in grey.
